# Allopatric divergence of cooperators confers cheating resistance and limits the effects of a defector mutation

**DOI:** 10.1101/2021.01.07.425765

**Authors:** Kaitlin A. Schaal, Yuen-Tsu Nicco Yu, Marie Vasse, Gregory J. Velicer

## Abstract

Social defectors may meet diverse cooperators. Genotype-by-genotype interactions may constrain the ranges of cooperators upon which particular defectors can cheat, limiting cheater spread. The bacterium Myxococcus xanthus undergoes cooperative multicellular development, but some developmental defectors cheat on cooperators during this process. In this study, interactions between a cheater disrupted at the signaling gene *csgA* and allopatrically diversified cooperators reveal a very small cheating range. Expectedly, the cheater failed to cheat on all natural-isolate cooperators owing to non-cheater-specific antagonisms. Surprisingly, lab-evolved cooperators that diverged from their cheating-susceptible ancestor by fewer than 20 mutations and without experiencing cheating had already exited the *csgA* mutant’s cheating range. Cooperators might also diversify in the potential for a mutation to reduce expression of cooperative trait or generate a cheating phenotype. A new *csgA* mutation constructed in several highly diverged cooperators generated diverse sporulation phenotypes, ranging from a complete defect to no defect, indicating that genetic backgrounds can limit the set of genomes in which a mutation creates a defector. Our results suggest that natural populations feature geographic mosaics of cooperators that have diversified in their susceptibility to particular cheaters and in the phenotypes generated by any given cooperation-gene mutation.

**Significance statement:** Selection on cooperators exploited by obligate cheaters can induce evolution of resistance to cheating. Here we show that cooperators can also rapidly evolve immunity to cheating simply as a byproduct of evolutionary divergence in environments in which cooperation and cheating at the focal trait do not occur because the trait is not expressed. We also find that differences in the genomic context in which a cooperation-gene mutation arises can profoundly alter its phenotypic effect and determine whether the mutation generates a social defect at all - a pre-requisite for obligate cheating. These findings suggest that general divergence of social populations under a broad range of environmental conditions can restrict both the set of mutations that might generate social defectors in the first place and the host range of such defectors once they arise.

---

Expressing a cooperative phenotype substantially less than conspecifics is often referred to as social defection (1). In microbes, such defection is often caused by mutations that intrinsically reduce expression of a cooperative trait, a type of defection we focus on here. One possible consequence of such defection is ‘cheating’ (1, 2), a social phenotype in which a defector gains a fitness advantage over cooperators by benefiting from their higher expression of a cooperative trait while not incurring its cost. However, it is possible that a defector may not gain such an advantage over all cooperative genotypes. Whether the defector genotype is able to display a cheating phenotype may depend on the social context; the ratio of defectors to cooperators (3) or the genotype of the cooperator may shape the nature (cheating or not) and strength of the interaction. In this scenario, it is then important to determine which cooperative genotypes within a diverse population are sufficiently compatible with a given defector to allow cheating upon interaction. This set of cooperators susceptible to cheating defines the ‘cheating range’ of that defector, analogous to a parasite’s host range (4–6). If a defector is unable to cheat on any cooperator, including its own cooperative parent (for whatever mechanistic reason), its cheating range is zero; it is never a cheater in any social context. For most microbial defectors that cheat at least on the parent from which they arose by mutation, the breadth of their cheating range is unknown.

Divergence among cooperators might impact not only the outcomes of social interactions between defectors and cooperators, but also the character of phenotypic effects of a given mutation as a function of the cooperator genotype. A particular mutation may result in the same degree of social defect or cause the same social-interaction phenotype between the mutant and its parent regardless of the cooperative background in which it appears. Alternatively, the mutation may be subject to genetic background effects that limit the degree to which it reduces a cooperative phenotype in certain genetic backgrounds, or even limits the set of cooperative genotypes within which it creates any defect at all. This may be referred to as the ‘defection-phenotype’ range of the mutation. When studying cheating phenotypes, it is important to consider the defection-phenotype range of the focal mutation when generalizing to the maintenance of cooperation within a system, in order to understand fitness effects of the mutation and whether a defective mutant will spread through a heterogenous population (see Fig. 1 for a conceptual overview).

**Figure 1.**
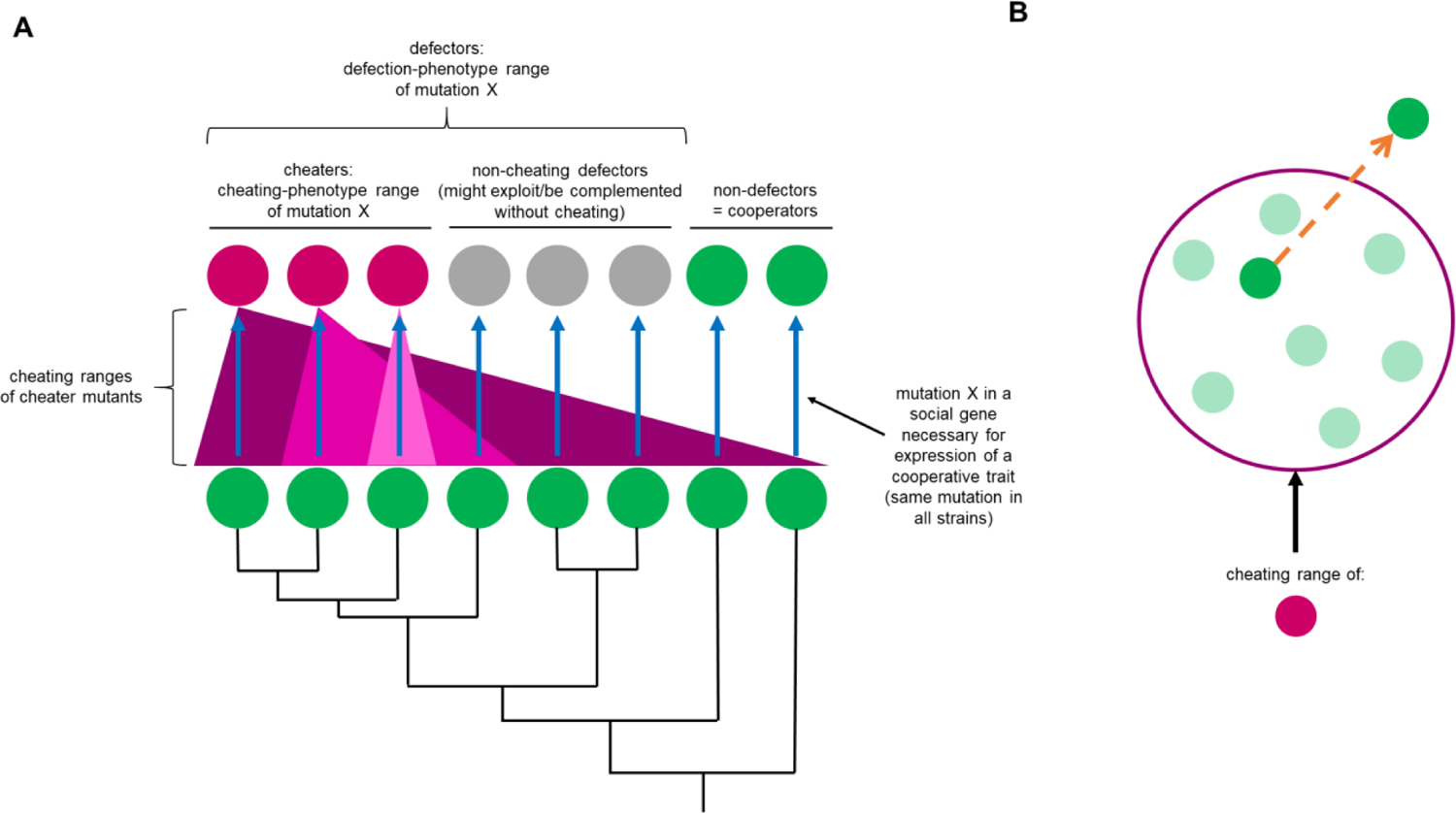
Concept visualization: cheating range and diverse possible effects of a defector mutation across genetic backgrounds. See *Semantics* in Methods for definition of cheating. (A) In nature, diverse cooperative genotypes (black phylogeny and green circles) may experience the same mutation, say mutation X, in a social gene (blue arrows). In some genetic backgrounds, mutation X may produce a cheater (magenta circles). Some cheaters may be able to cheat on many different cooperative genotypes, i.e., they have a wide cheating range (darkest pink triangle). Other cheaters may cheat only on their cooperative parent (and likely nearly-identical genotypes), i.e., they have a narrow cheating range (lightest pink triangle). In other genetic backgrounds, mutants with mutation X may not able to cheat even on their own cooperativ e parents and may be referred to as non-cheating defectors (grey circles). Together, the cheaters and the non-cheating defectors represent the defection-phenotype range of mutation X. It is also possible that the mutation does not alter the cooperative phenotype enough to produce a defector (“non-defectors” green circles). B) Evolutionary change may cause a lineage of cooperators (orange arrow) to exit a given cheater genotype’s cheating range (dark pink ring). Such a transition might result from selection that targets cooperator fitness during cooperator-cheater interactions or, as highlighted in this study, other forces such as drift or selection unrelated to cheating.

Many mechanisms are known which can limit cheaters (e.g. (3, 7, 16–20, 8–15)). These mechanisms either passively or actively stabilize cooperative relationships by causing the benefits of cooperation to be directed more to cooperators than to cheaters, on average (7). Some of these mechanisms can evolve in cooperative systems in response to different types of selective pressures. This has been demonstrated in experiments with microbes, in which cooperators evolved to outcompete cheating defectors by adapting either to the cheaters themselves or to abiotic features of the environment in which cooperative and cheating phenotypes were expressed (12, 14, 17, 19, 21–23). In those studies, the relevant cooperative trait was important to fitness in the environments in which the cooperators were evolving, and the cheaters that exploited the trait were present during evolution. It may be possible, however, for cooperators to latently evolve resistance to being cheated on, while adapting to an environment in which the relevant cooperative trait is not even expressed. In other words, a cooperative genotype may be shifted outside of a given defector’s cheating range by evolution unrelated to cooperation and cheating and which occurs in allopatry from the defector.

Allopatric divergence often profoundly alters biotic interactions. For example, allopatric speciation is an important form of divergence in animals and plants in which barriers to reproduction evolve between spatially separated populations (24, 25). In social insects, allopatric divergence affects social-parasitism behaviors (26, 27). In microbes, allopatric divergence of social types can increase inter-strain antagonism (28) and also generate kin-discriminatory colony interactions (29), fitness asymmetries specific to social interactions (30), and social exploitation among cooperation-proficient genotypes (30). However, the potential for allopatric divergence between cooperators and defectors to constrain the cheating ranges of defector genotypes remains little explored.

The soil bacterium *Myxococcus xanthus* is a well-studied microbial cooperative system that displays social behaviors such as swarming motility (31) and multicellular development into spore-filled fruiting bodies (32). Cooperative lab strains of *M. xanthus* that are proficient at development sometimes yield obligate defector mutants, genotypes that constitutively produce less of a functional signal molecule necessary for normal development (3). Some such defectors can cheat on the cooperator from which they immediately descend – that is, the mutant has higher fitness than the cooperative parent in mixed groups due to the mutation that causes its cooperation defect. Lab-derived cheaters resulting from evolution experiments (3) or mutagenesis (33, 34) can cause major population collapses due to cheating load (35–37), sometimes driving entire populations they inhabit to extinction (8). Analogous social collapse due to conspecific social parasitism has been documented in the African honeybee (38). Cheating defectors that emerge in nature therefore have the theoretical potential to devastate the populations in which they arise.

Natural *M. xanthus* populations living in spatially-structured soil environments have high levels of genetic diversity even at small scales (28, 39). Such diversity includes positively frequency-dependent antagonisms directed broadly against many conspecifics (39) that occur pervasively among developmentally proficient natural isolates. These antagonisms are often lethal, and may be expressed during both vegetative growth and starvation-induced development, with minority genotypes almost always losing to majority genotypes irrespective of fitness outcomes in 1:1 mixes. Such antagonisms are predicted to be a major determinant of cheating ranges in nature, perhaps rendering most developmentally-proficient cooperator genotypes unsusceptible to cheating to most cheaters derived by mutation of diverged cooperators (39).

Several cheater genotypes have been studied in *M. xanthus* (3, 40). For example, mutations in the genes *asgB* (41) and *csgA* (34) prevent mutants from producing signal molecules (A-signal and C-signal, respectively) which are necessary for the early stages of fruiting body formation in the type strain DK1622. Both mutations create obligate social defectors by reducing spore production in clonal groups by several orders of magnitude (3). Here we focus on the *M. xanthus* cheater DK5208 (also known as LS523, see Methods), which has a transposon insertion in *csgA* (34). While this gene is necessary for normal development in the type strain, the precise mechanisms of C-signaling are debated. Earlier research suggested that the C-signal is a 17-KD fragment of CsgA which acts as an outer membrane signal and impacts developmental timing (42–44), while more recent studies suggest that the signal may derive from lipids generated by CsgA phospholipase activity in starving cells (45). To date, cheating phenotypes of *M. xanthus* defectors have been studied primarily in the social context of pairwise interactions with their cooperative parent or a recent ancestor.

Here we first investigate the cheating range of the *M. xanthus csgA* mutant DK5208, and in particular the role of allopatric divergence (as opposed to defector-induced evolution) in shaping this range. We test the ability of the defector, which cheats on its parent, to cheat on natural strains which we expect diverged from it in allopatry, as they were isolated at large geographic distances from each other and from the original isolation site of DK5208’s ancestor. We examine whether known allopatric divergence generated *de novo* in the laboratory, in environments in which the relevant cooperative trait is not expressed, can latently generate barriers to social exploitation (*i.e.* shift the evolved strains out of DK5208’s cheating range) by testing the cheater’s ability to exploit closely-related descendants of its cooperative parent strain. We then test for a different kind of potential consequence of allopatric divergence among cooperators, namely genetic-background effects (46, 47) on the social phenotype caused by a mutation in a cooperation gene. Specifically, we test whether disrupting *csgA* in the same manner in several allopatrically diverged cooperator natural isolates results in different social phenotypes as a function of cooperator genomic background. Collectively, our results suggest that allopatric divergence of both the genomic contexts in which mutations in social genes arise and of the social contexts in which resulting mutants might interact play greater roles in determining the evolutionary fates of non-cooperation alleles than has been previously appreciated.

## Materials and Methods

### Semantics

In this study, we use ‘obligate cheating’ (or simply ‘cheating’) to refer to a social interaction in which one interactant (the cheater) is obligately defective to some extent at expressing a focal cooperative trait relative to a cooperative genotype, and yet gains a relative fitness advantage over the cooperator by social exploitation when they interact under relevant conditions (e.g. (8, 10)). Here we do not consider strains that are intrinsically proficient at a high level of cooperation yet outcompete other cooperation-proficient strains in mixed groups during a cooperative process (which are sometimes called ‘facultative cheaters’ (9) or ‘facultative exploiters’ (30)), as their competitive success does not undermine the persistence of cooperation *per se*. We use ‘social exploitation’ to refer to any social interaction from which a focal partner derives an absolute-fitness benefit, even if the other interactant(s) are not harmed (48).

### Strains and growth conditions

As the defector, we used the *M. xanthus* developmental mutant DK5208 (Table S1), which is a yellow clonal isolate of the strain LS523 (17, 49). LS523 is a mutant of the wild-type strain DK1622 containing a Tn5 transposon insertion in the *csgA* gene (position 217 out of 690 bp) which confers resistance to oxytetracycline (34, 50). We used a diverse set of developmentally-proficient cooperative *M. xanthus* strains: (i) a wild-type laboratory strain and a rifampicin-resistant derivative strain, (ii) thirteen natural isolates from around the world, (iii) ten natural isolates from Bloomington, Indiana, USA (the latter two sets of strains referred to as ‘N’ for ‘natural isolate’), and (iv) nine experimentally-evolved clones (referred to as ‘E’ for ‘evolved’) that descended from the wild-type strain during a previous evolution experiment and retained high levels of spore production (Table S1).

The wild-type laboratory strain GJV1 (strain ‘S’ in (51)) is derived from DK1622 via a small degree of sub-culturing and differs from it by five mutations (52). DK1622 (GenBank accession number CP000113) is a mutant of the natural isolate DK101 (also called FB) which was constructed to restore full function to a motility system that had acquired a mutation during the laboratory culturing process following isolation (53, 54). The sample of *M. xanthus* strain FB referred to as DK101 was obtained from the University of California, Berkeley culture collection (53, 55, 56). Strain FB (ATCC 25232) was sourced from strain Beebe 1941 (ATCC 19368), which was isolated from soil in the vicinity of Ames, Iowa around 1941 (57, 58) and which has since been lost, making FB the new ancestral laboratory strain for this species. However, DK1622 is the most commonly used wild-type strain. For a summary of the culturing history of these strains, from the Beebe isolate to DK1622, see Fig 5 in ref. (59). As the lineages resulting in the samples of GJV1 and DK5208 used in this study have been maintained in the lab since the original isolation event in Ames, Iowa in 1941, while the natural isolates we used (iii and iv) were isolated after the year 2000 from multiple sites around the world, the closest being Bloomington, Indiana (828 km away), we assume that there has been allopatric divergence between GJV1/DK5208 and the natural isolates. In experiments testing whether the effects we observed were specific to the defector strain, we used GJV2, a spontaneous rifampicin-resistant mutant of GJV1 (aka strain ‘R’ in (51), see also (12, 60)).

We selected the natural isolates from previously-published analyses of *M. xanthus* relatedness in nature based on collections either of natural isolates from around the world (61) or of natural isolates taken from carefully-defined distances in Bloomington, Indiana, USA (62). We chose strains that grew well and sporulated proficiently under laboratory conditions. To allow for the possibility of differences in interactions with DK5208 based on geography or relatedness at this scale, we chose pairs of strains that were isolated from the same location (61) or at known centimeter-, meter-, and kilometer-scale distances from each other (62) when growth patterns permitted.

The evolved clones come from an evolution experiment now referred to as MyxoEE-3 (63) and were isolated from populations descending from either GJV1 or GJV2 and which had undergone 40 two-week cycles of evolution as motile colonies expanding on 0.5% or 1.5% nutrient agar, an environment in which development does not happen, as described previously (29, 30, 64). In this case, the allopatric divergence between GJV1/DK5208 and the MyxoEE-3 clones occurred under controlled laboratory conditions for a known period of time.

We inoculated frozen *M. xanthus* stocks onto CTT (65) 1.5% agar plates and incubated at 32 °C and 90% rH for 4-5 days, after which we inoculated colony samples into CTT liquid and grew them overnight (32 °C, 300 rpm). Where appropriate, we supplemented media with 40 μg/ml kanamycin or 12.5 μg/ml oxytetracycline.

### Plasmids and mutant construction

We disrupted *csgA* (NCBI DK1622 locus tag MXAN_RS06255, old locus tag MXAN_1294) at the same nucleotide position in *M. xanthus* strains GJV1, Chihaya 20, Serengeti 01, GH3.5.6c2, and MC3.5.9c15 by plasmid insertion, without otherwise altering the native *csgA* sequence of each strain. We transformed each strain with a plasmid containing a fragment of the strain’s own *csgA* allele. We amplified all *csgA* fragments for the different strains from the identical region of *csgA* using the same two primers, which pair to fully conserved *csgA* segments (see Fig. S3A). We selected PCR primer sequences in the conserved *csgA* segments by aligning the *csgA* sequence of the published *M. xanthus* DK1622 genome (NCBI:txid246197) with natural-isolate genomes (SRA accession numbers: SRR8298023, SRR8298022 (61, 66); Fig. S3A). Phylogenetic relationships among the *csgA* sequences are shown in Fig. S3B. PCR-amplification using forward primer 5’-<u>T</u>AATTCGTCCAGCAGCTCCTGCTGC-3’ (*csgA* positions 44-67, genome positions 1520242-1520265 (underline indicates point mutation introduced to disrupt restriction enzyme site)), and reverse primer 5’-<u>TTA</u>CCCATCCGCGAGGTGACGTG-3’ (*csgA* positions 394-413, genome positions 1520592-1520611 (underline indicates added stop codon)) resulted in a 370-bp internal fragment of *csgA* from each of the natural isolates and GJV1. We purified the fragments using the QIAquick^®^ PCR Purification Kit (QIAGEN, Hilden, Germany) and verified their length on a 1% agarose gel, then ligated each insert into the pCR-Blunt vector (Invitrogen, San Diego, CA), which carries a kanamycin resistance marker. We verified the pCR-*csgA*413 plasmids (Table S3) bearing the individual *csgA* fragments by sequencing (Microsynth AG, Balgach, Switzerland). We electroporated each *M. xanthus* strain with the respective plasmid to create merodiploids with truncated copies of *csgA*. The CsgA protein encoded by the first partial *csgA* sequence is truncated at its amino acid position 138 by the plasmid integration. The second copy is deprived of the native *csgA* promoter and its 5’ terminal sequence contains a stop codon engineered at the beginning of the amplified *csgA* fragment. Thus, successful transformants are designed to produce a truncated CsgA protein 137 aa long, or ~60% of the full-length 229-aa GJV1 CsgA protein. We verified transformants by diagnostic PCR and antibiotic-resistance phenotype.

### Developmental assays

We centrifuged exponential-phase liquid cultures (5000 rpm, 15’) and resuspended in nutrient-free TPM (67) pH 8.0 to a density of ~5 × 10^9^ cells/ml. We inoculated 50 μl of each strain or mixture onto TPM pH 8.0 1.5% agar plates to initiate development and incubated for 72 hours (32 °C, 90% rH). We made all mixes at a 1:99 ratio by mixing 1 μl of the *csgA* mutant and 99 μl of the cooperator. We harvested entire starved populations with sterile scalpels and incubated each sample in 1 ml of ddH_2_O for 2 hours at 50 °C to kill any non-spore cells, then sonicated to disperse spores. We dilution-plated the samples into CTT 0.5% agar to count the number of CFUs. For mixed competitions, we plated the samples into agar with and without antibiotic to generate counts for the *csgA* mutants and total population, respectively, and therefore by subtraction counts for the cooperators. We plated pure-culture samples of *csgA* mutants with antibiotic.

### Data analysis

We performed all data analysis and statistical testing using R version 4.0.0 and RStudio version 1.2.5042. We visualized the data using the ggplot2 package (68). Original data files and analysis protocols, including statistical scripts, R Markdown files, and full results of statistical tests, may be accessed via Dryad (doi: 10.5061/dryad.fbg79cnsb). For the *csgA* mutants in the developmental assays, we calculated the one-way mixing effect

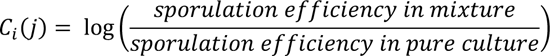

where “sporulation efficiency” refers to the fraction of cells inoculated that became spores, and the relative fitness in mixture

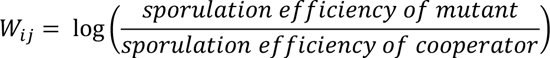

as in Vos and Velicer (28). Here, ‘*i*’ refers to the mutant and ‘*j*’ refers to the cooperator. We analyzed our experimental results using general linear models followed by post-hoc Tukey HSD tests, Dunnett tests, or *t*-tests, as reported in the results. For calculations of *C_i_(j)* and *W_ij_*, we assumed that strains for which we counted zero spores produced the maximum number of spores which would have been below the detection limit of our dilution plates.

We constructed the phylogeny in Figure S3B using Clustal Omega version 1.2.1 (69), PhyML version 3.1 (70), and Newick Display (71) via the Galaxy platform (galaxy.pasteur.fr; (72)).

## Results

### Cheating range of a defector strain

The M. xanthus social defector strain DK5208 is known to cheat on the developmentally proficient lab strain GJV1 during starvation. DK5208 converts a greater proportion of its vegetative cells into spores than does GJV1 in these mixed groups, despite producing far fewer spores than GJV1 in monoculture (3). The two strains are closely related; GJV1 is a sub-cultured recent descendent of DK1622 (52), the strain from which DK5208 was created (see Strains and growth conditions in Materials and Methods). It has been hypothesized that inter-strain antagonisms may restrict the set of cooperative genotypes on which a given defector can cheat (39), leading us to hypothesize that genetic divergence may be a factor shaping the cheating range of such a defector.

#### Cheating range excludes distantly related natural isolates

To test the effect of high degrees of genetic divergence on cheating range, we mixed DK5208 in the minority (1:99) with a diverse set of natural strains (ii and iii in *Strains and growth conditions* in Materials and Methods) isolated from various locations around the world (61, 62) and allowed the pairs to interact during starvation. We compared spore production by DK5208 in mixes with GJV1 versus with the 23 natural isolates, which we expect to differ from GJV1 by at least tens of thousands of mutations (see Fig. 1 in ref. (61), Fig. S2 in ref. (66)). As expected, DK5208 cheated on GJV1 (*p* = 0.005, one-sided *t*-test for *Wij* > 0). However, not only did DK5208 fail to cheat on any of the natural isolates (mean *Wij* values < 0, *p*-values < 0.005, 23 two-sided *t*-tests against 0 with Bonferroni-Holm correction; Fig. 2), we detected zero or extremely few spores in all pairings (detection limit = 10 spores; Fig. S1A).

**Figure 2.**
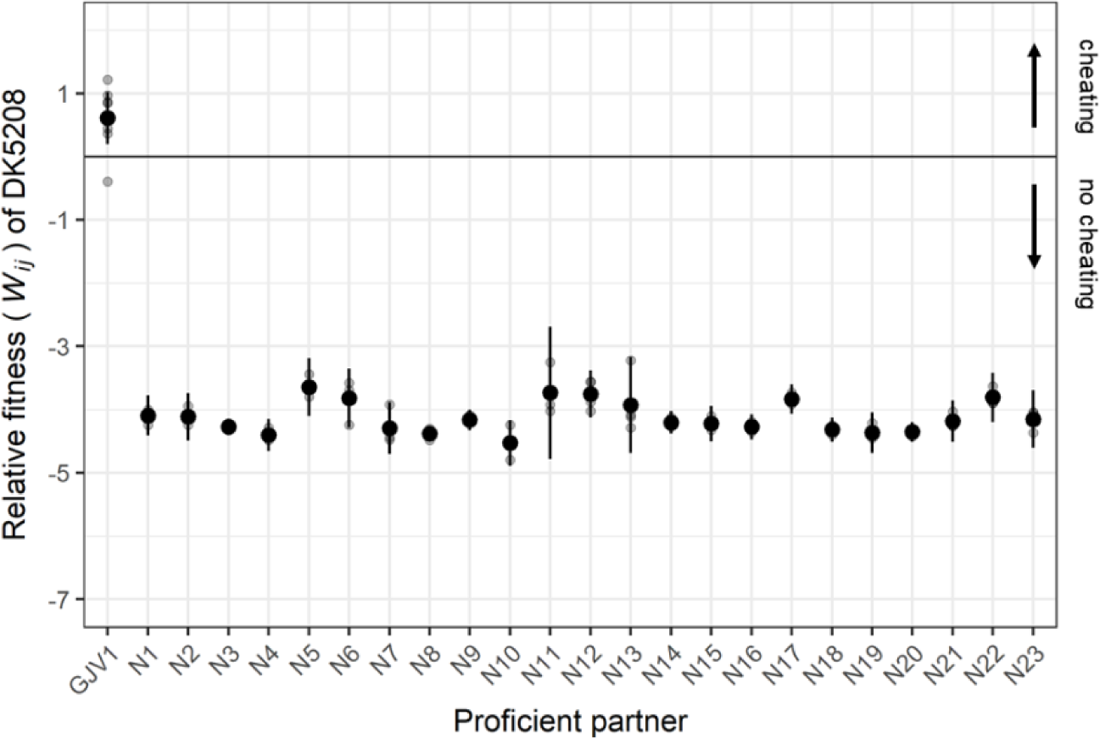
DK5208 cheats on a closely related cooperator but is outcompeted by distantly related natural isolates. Sporulation fitness (*W_ij_*) of DK5208 relative to GJV1 and 23 natural isolates in initially 1:99 (DK5208:competitor) mixed cultures. Grey circles represent individual replicate estimates, black circles show cross-replicate averages, and error bars show 95% confidence intervals; 3 or 4 biological replicates for natural isolate mixes, 8 for GJV1.

To test whether defector-independent interactions, rather than defector-specific mechanisms, were responsible for preventing DK5208 from sporulating in the preceding experiment, we mixed a developmentally proficient, rifampicin-resistant variant of GJV1 (GJV2) with the natural isolates (Fig. S1B). Although we observed a greater negative marker effect on GJV2 sporulation than observed in prior studies (30, 51), GJV2 nonetheless produced substantial numbers of spores in mixture with GJV1. However, like DK5208, it produced no detectable spores when mixed with the natural isolates. The failure of GJV2 to sporulate in these mixtures indicates that mechanisms not specifically targeting defectors prevent DK5208 from cheating on these strains, thereby placing them outside of DK5208’s cheating range.

#### Small degrees of allopatric divergence can eliminate cheating upon secondary contact

Since DK5208 was unable to sporulate at all in mixtures with genetically distant natural isolates, we tested whether smaller degrees of divergence might also reduce or eliminate DK5208’s ability to cheat. We performed additional developmental competitions using developmentally-proficient clones from nine experimental populations (iv in *Strains and growth conditions* in Materials and Methods) that descended from GJV1 in an evolution experiment recently named MyxoEE-3 (63, 64). The MyxoEE-3 clones examined here evolved in nutrient-rich environments in which starvation-induced cooperative development was not expressed. They were therefore not under selection to improve fitness during development. These populations also evolved in the absence of DK5208 (or any other known cheater) and therefore had no opportunity to interact in an evolutionarily relevant way with any genotypes capable of developmental cheating (64). Clones isolated from these MyxoEE-3 populations had each accumulated no more than 20 mutations (64). As expected from previous experiments, DK5208 cheated on GJV1 (Figs. 3 and S2; *p* = 0.016, one-sided *t*-test for *Wij* > 0). However, the evolved MyxoEE-3 clones exhibited a clear trend of decreased susceptibility to cheating. DK5208 relative-fitness estimates were lower against eight of the nine evolved clones than against GJV1 (all except E6, Fig. 3), an outcome unlikely to have occurred by chance (one-tailed sign test, *p* = 0.02).

**Figure 3.**
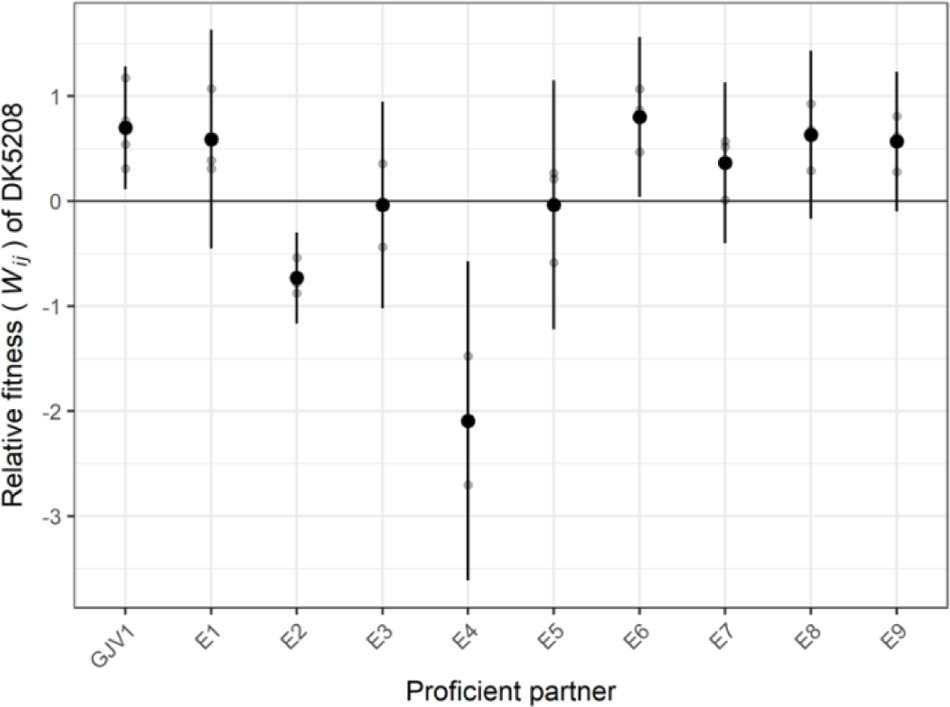
DK5208 cheats on GJV1 but not on some of GJV1’s closely related descendants. Sporulation fitness (*W_ij_*) of DK5208 relative to GJV1 and descendants of GJV1 from MyxoEE-3 (*W_ij_*) in initially 1:99 (DK5208:competitor) mixed cultures. Symbols as in Fig. 1; 3-4 biological replicates.

DK5208 had much lower fitness against two evolved clones in particular – E2 and E4 – than against the ancestor GJV1 (*p*-values < 0.001, Dunnett test for difference from GJV1). DK5208 not only failed to outcompete E2 and E4 (Figs. 3 and S2; *p*-values > 0.1, 9 two-sided *t*-tests against 0 with Bonferroni-Holm correction) but in fact appears to be outcompeted by them, as it was by the natural isolates (mean and 95% confidence interval of *Wij* = −0.7 [−1.2, −0.3] and −2.1 [−3.6, −0.6], respectively). Thus, the ten mutational steps each which separate E2 and E4 from GJV1 (Table S2) are enough to alter the fitness ranks emerging from the social interaction, eliminating the cheating phenotype. This illustrates that even a small degree of evolution in an environment in which a focal cooperative trait is not expressed can latently generate resistance to cheating in environments in which cooperation does occur. This small degree of allopatric, cheater-blind evolution has already shifted some strains, E2 and E4, out of DK5208’s cheating range and we predict further evolution would do the same for other strains.

### Defection-phenotype range of a mutation

Having shown that DK5208 has a narrow cheating range, beyond which some cooperators can evolve by only a few mutations separating them from the cheater’s parent, we then sought to test the defection-phenotype range of a mutation in *csgA*. We hypothesized that mutations in *csgA* would create a similar phenotype (a developmental defector with a narrow cheating range) in the set of allopatrically diverged genetic backgrounds that we tested.

#### Social defection phenotypes are subject to genetic-background effects on csgA disruption

We selected four of the natural isolates used in the assay reported in Fig. 2 and introduced identical disruptions of their native *csgA* alleles. As we used a different method from that used to construct LS523 (DK5208’s direct ancestor), we included GJV1 as a control. We constructed plasmids which integrated into each strain at *csgA* by amplifying a 370-bp internal fragment of each strain’s allele. We used primer sequences within *csgA* that are fully conserved across all strains (Fig. S3A) and ligated each fragment into the pCR-Blunt plasmid vector (Table S3). Transformation with the resulting plasmids successfully disrupted *csgA* at base-pair position 413 (out of 690 bp, see Methods) in all five strains and created partial-copy merodiploids at that locus. Thus, we inserted an identical plasmid vector sequence into *csgA* in each strain without otherwise altering its native *csgA* sequence. *A priori*, we anticipated that this mutation would cause pure-culture sporulation defects in all strains and would likely create a cheating phenotype in at least the GJV1 background, as well as possibly other backgrounds. But at the same time we also anticipated that quantitative effects of the mutation on both pure-culture sporulation and sporulation of the mutant when mixed with its parent might vary, with one possible outcome being that the mutation might create a cheating phenotype in some backgrounds but not others.

Indeed, the five resulting *csgA* mutants vary greatly in their monoculture developmental phenotypes despite carrying the same plasmid-insertion mutation, indicating that phenotypic expression of this mutation is subject to genetic-background effects. All four natural-isolate mutants produced far fewer spores than their parental strains in monoculture (max. ~1% of parent; *p* < 0.032 for the four natural isolates, five one-sided *t*-tests against 0 with Bonferroni-Holm correction; Fig. 4). The four natural-isolate mutants (N9 *csgA*, N16 *csgA*, and N23 *csgA*) produced fewer spores than their parents, and N2 *csgA* produced no detectable spores (detection limit = 10 spores; Fig. S4). However, the mutation had no significant effect on spore production in the GJV1 background (*p* = 0.26, same *t*-tests as above) and thus did not create a developmental defector. The plasmid insertion is in a different position in GJV1-csgA than the transposon insertion in DK5208 (position 413 vs 217), so we performed a further test which showed that the position of plasmid insertion within GJV1’s *csgA* matters to an extent in terms of the resulting social-defection phenotype (Fig. S6), but the pattern and the connection to CsgA function are unclear and require further investigation. We found no evidence that disrupting *csgA* at a different position, closer to position 217, would have created a cheating phenotype similar to that of DK5208 (Fig. S6).

**Figure 4.**
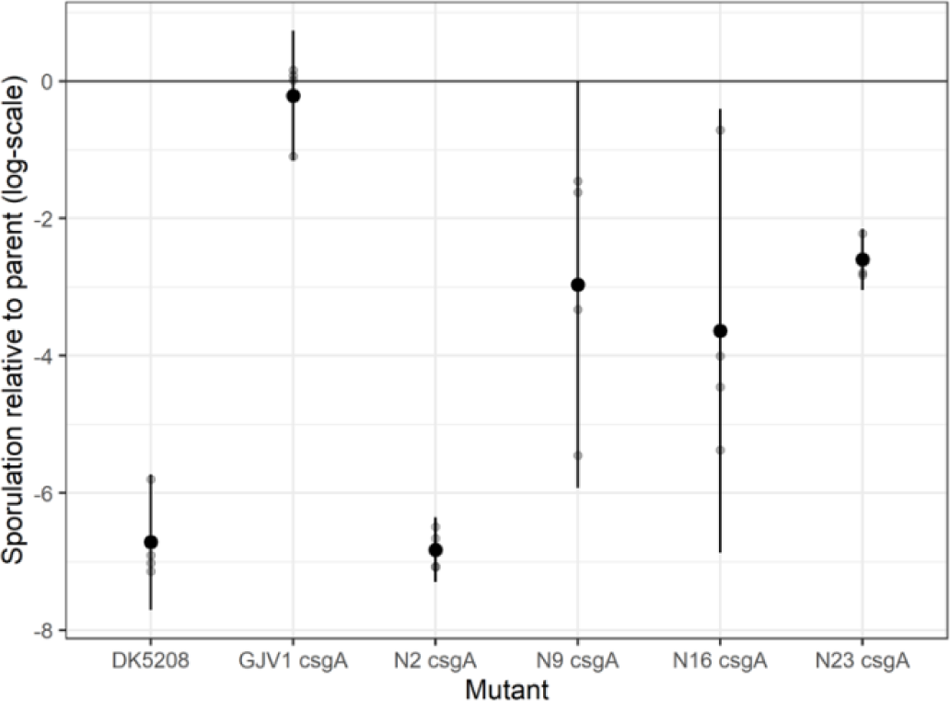
Genetic background effects on the phenotype of a *csgA* mutation. We show the pure-culture spore production of each new *csgA* mutant relative to that of its parent. The DK5208 *csgA* transposon mutation severely reduces spore production relative to GJV1, whereas the new *csgA* plasmid-disruption mutation in the GJV1 background does not. Genetic background effects: The same *csgA* plasmid disruption impacts relative spore production very differently depending on the parental genotype, thus revealing strong epistatic interactions and developmental system drift. Symbols as in Fig. 1; 4 biological replicates.

These outcomes indicate that different genomic backgrounds require different lengths of uninterrupted *csgA* to achieve high levels of sporulation; 413 base pairs of the GJV1 *csgA* allele were sufficient in the GJV1 background, but the same length allele was insufficient in the other backgrounds, especially in N2. Thus, we find that an identical mutation can have different effects on a social phenotype depending on the genomic context in which it occurs. Allopatric divergence among cooperators altered the genetic requirements for expression of a cooperation-based trait – in this case, the length of intact *csgA* gene required to allow high levels of spore production.

Because GJV1-*csgA* is not a defector, in this system it does not have the potential to produce a cheating phenotype when interacting with a compatible cooperator. However, since the other mutants exhibited significant defects, we could test whether they cheated on their parent strains. All four defectors were fully complemented by their own parental strains in mixed groups (Figs. 5, S4 and S5), but we detected no significant differences in relative fitness for any of the mutant-parent strain pairs (*p*-values > 0.05, 26 two-sided *t*-tests against 0 with Bonferroni-Holm correction; Fig. 5). In other words, the defectors socially exploited their parents for gains in absolute fitness, but these gains were not high enough to constitute cheating.

**Figure 5.**
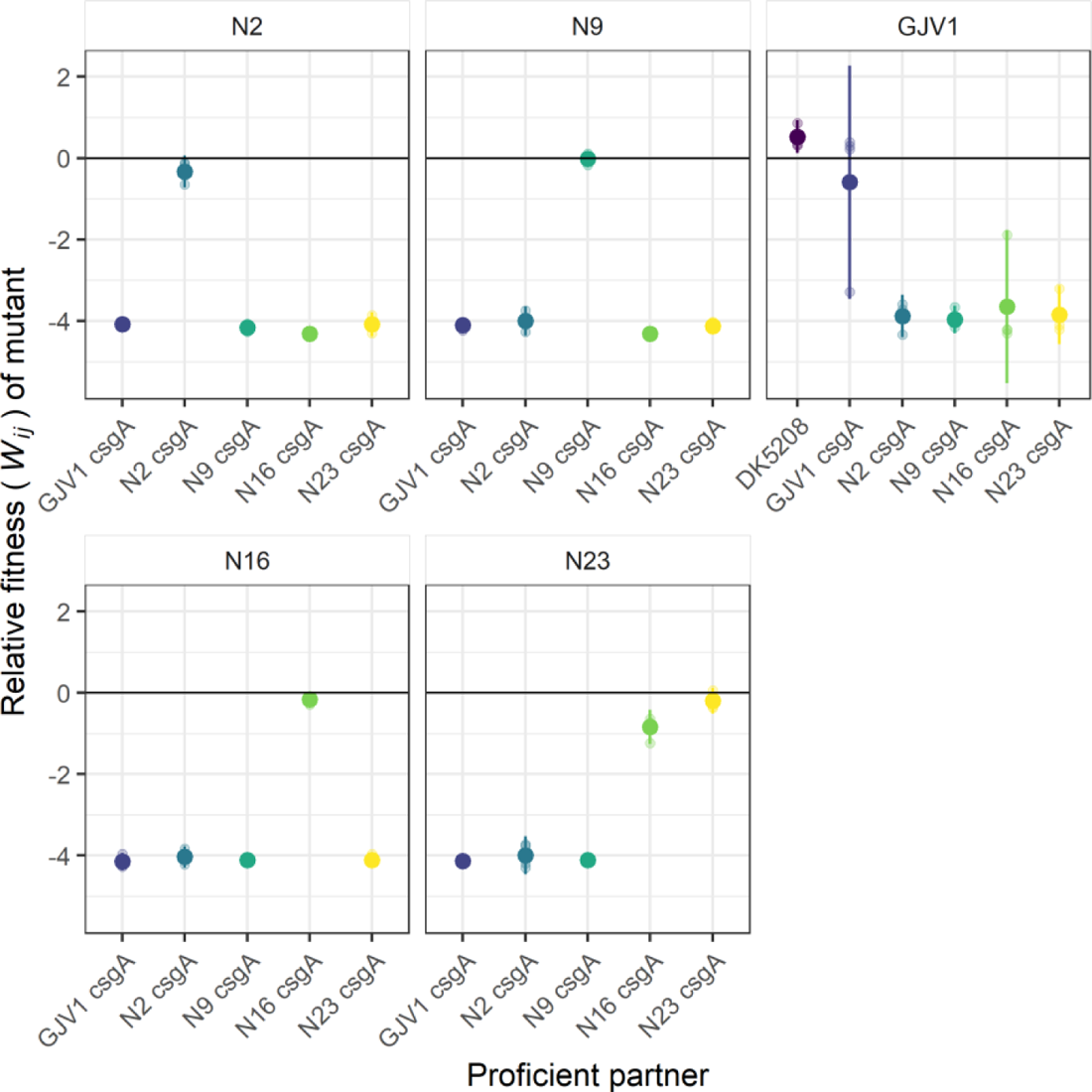
New *csgA* mutants have relative fitness similar to their parent in mixed groups but are generally outcompeted by non-parental cooperators. Sporulation fitness (*W_ij_*) of *csgA* mutants relative to developmentally proficient partners in initially 1:99 (mutant:partner) mixed groups. Symbols as in Fig. 1; 4 biological replicates.

We then tested how well these mutants would fare in mixture with the other three natural isolate parents and GJV1. Just as DK5208 was unable to exploit any natural isolate for gains in absolute fitness (Fig. S1A), the plasmid-disruption *csgA* mutants were generally not complemented in mixes with cooperation-proficient strains other than their own parent. Indeed, in most cases, the mutants produced either no spores or extremely few in mixture with nonparental strains (Fig. S4). One surprising exception was the mixture of N16 *csgA* with N23, in which the mutant was complemented but was also not significantly harmed (*p* = 0.51, 26 two-sided *t*-tests against 0 with Bonferroni-Holm correction). This suggests that, independent of the *csgA* mutation, the N16 background has high enough fitness relative to N23 to compensate for being in the minority in mixture.

## Discussion

Cheating of social defectors on cooperative types is well-studied but is in some respects still poorly understood. Building on the idea that manifestation of cheating can be dependent on social context (frequency and identity of interacting partners, e.g. refs. (3, 73)), we describe social defectors as having a ‘cheating range,’ or a set of cooperative genotypes upon which they can cheat, *i.e.* socially exploit to gain a relative fitness advantage, similar to the host ranges of parasites. Here, to demonstrate the cheating range concept, we examined the role of allopatric divergence in the expression of cheating phenotypes in *M. xanthus* social defectors, as well as the possibility of genetic-background effects to limit the set of genotypes in which a given mutation creates a social defector capable of cheating.

To demonstrate the cheating range concept and explore the role of allopatric divergence in moving cooperators beyond a given cheater’s cheating range, we allowed the cheater mutant DK5208 to interact with two sets of cooperator genotypes that were either distantly or closely related the cheater. Although DK5208 cheats on a strain (GJV1) that is nearly identical to its parent (DK1622), it not only fails to cheat on genetically distant natural isolates (Fig. 1) but effectively fails to sporulate at all (Fig. S1A). The extremely low fitness of this defector against the natural isolates appears unrelated to the developmental defect that allows it to cheat on GJV1. Sporulation by two different sporulation-proficient mutants of GJV1 – GJV2 and GJV1 *csgA* – is also effectively eliminated by the same natural isolates when they interact (Figs. S1B and S4). Toxin-mediated antagonism between natural isolates often occurs during both growth and development (39), and these inter-strain antagonisms clearly contribute to placing the natural isolates outside of DK5208’s cheating range.

Due to the distant sampling locations of the relevant strains and our understanding of *M. xanthus* biogeography (see *Methods, Strains and growth conditions*), we assume that the natural isolates evolved for many generations in allopatry from the natural isolate from which the defector is derived, but the detailed evolutionary histories are in fact unknown and the total sets of genetic differences that prevent DK5208 from cheating on these natural isolates could be very difficult to comprehensively identify. So we allowed DK5208 to interact with lab-evolved cooperators which were very closely related and whose evolutionary history since divergence was known. We asked how readily social barriers to cheating might evolve in cooperative lineages in environments in which they do not even express a cheatable cooperative trait and do not meet defectors at all. That defector resistance might emerge in such a fashion was plausible in light of other novel social interactions known to evolve indirectly, including cheating by mechanistically obligate defectors (3), facultative social exploitation (30), and kin discrimination in the form of colony-merger incompatibilities (29).

When mixed with clones from the MyxoEE-3 evolution experiment (closely related lab-evolved cooperators that had no history of experimental selection on development because they evolved in nutrient-rich environments and in the absence of the defector), DK5208 exhibited a trend of reduced fitness compared to mixture with the experimental ancestor, and it failed to cheat on multiple evolved cooperators (Fig. 2). This outcome was not obvious, as we might have expected the evolved strains to have actually decreased in fitness relative to the defector during MyxoEE-3 due to the combination of a lack of selection on traits related to starvation and the presence of selection for adaptation to resource abundance. Yet even this brief allopatric evolution in a nutrient-rich environment was sufficient to begin to shift these cooperators toward the edge of and beyond DK5208’s cheating range. This suggests that for *M. xanthus* social defectors, the cheating range should be explored at the scale of 10s of mutations, rather than 100s or 1000s (Table S1). It also suggests that, as genetic diversity is high even at small spatial scales (66, 74, 75), the close proximity of noncompatible cooperators may often greatly limit the spatial spread of any given cheater genotype.

Just as lineages of cooperators can shift outside a cheating range due to diversification,, so too might such diversification actually limit the range of genotypes in which a given mutation creates a defector or cheater in the first place, due to genetic background effects (GBEs). Fruiting body development and sporulation in *M. xanthus* are complex processes involving many genes and regulatory pathways (31, 67, 76, 77). Evolutionary change affecting genes that contribute to a complex developmental process can alter their epistatic relationships, causing a given allele to produce different phenotypes in different genetic backgrounds (46, 47). GBEs have been found in many systems, including insects (78, 79), maize (80), mice (81), and microbes (82, 83). We tested for GBEs on the cheating phenotype we considered here by disrupting the developmental signaling gene *csgA* in a set of well-diverged cooperators, strains that produce similarly high numbers of spores. This disruption produced highly variable spore-production phenotypes – in one case (N2) effectively eliminating sporulation, while maintaining a wild-type sporulation level in the GJV1 background. These strains have thus diverged in how their genetic backgrounds epistatically interact with a plasmid-disruption mutation that renders the terminal portion of *csgA* unusable. The 5’ region of *csgA* left intact upstream of the disruption provides sufficient function to allow normal sporulation in the GJV1 background but not in the natural isolates. Genomic divergence between N2 and GJV1 altered the length of intact *csgA* necessary for cooperative spore production.

This outcome has both evo-devo and social-evolution implications. In aggregative multicellular systems, such as *M. xanthus*, evo-devo (84, 85) and social evolution are intrinsically intertwined because development is a cooperative process among reproductively autonomous organisms. From an evo-devo perspective, our *csgA*-disruption results demonstrate developmental system drift (DSD; (86)) among conspecifics in a microbial system. In DSD, the genetic basis of a developmental phenotype diverges across lineages, either stochastically (86) or due to selection (46, 87), while the fundamental phenotype itself is conserved. This phenomenon has been documented in a wide range of systems (88–90). In the myxobacteria, the gene sets necessary for fruiting body development have diverged extensively across species (91, 92). At much shorter evolutionary timescales, analyses of an experimental lineage (93) and the natural variation at a regulatory region (61) have shown that the genetic pathways underlying *M. xanthus* development are evolutionarily malleable (94). Our *csgA*-disruption results show that the general epistatic environment of these pathways can diverge sufficiently within the same species to render a gene or part of a gene conditionally essential to a major developmental phenotype (95).

From a social evolution perspective, here we show that diversification has generated epistatic effects on the phenotype resulting from disruption of a key cooperation gene, to the extent that elements of the gene necessary for normal development in some strains are not similarly necessary in all strains. We produced identical disruptions of *csgA* in diverged genomic backgrounds, demonstrating that whether a mutation generates a social defect, and therefore potentiates cheating, can depend on the genomic context. This suggests that different *M. xanthus* strains may differ in the sets of mutations that confer a cheating phenotype, which could limit the ability of any given cheating mutation arising in one genomic background to spread across genomic backgrounds by horizontal gene transfer (96, 97). Thus, in aggregatively multicellular microbes, just as social selection can drive evolution of developmental features (20, 93, 98), so too may divergence of developmental genetic architecture reciprocally shape social evolution.

Together, these outcomes suggest that cheaters arising in spatially structured natural populations (39, 62, 66, 74, 75) are likely to have cheating ranges that are narrow both genetically and, because genetic and spatial distance correlate in wild populations (99), geographically. Cheater-blind allopatric divergence, emerging due to drift or selection, may generate a patchy phylogeographic mosaic of cheater-cooperator compatibility types across which most cooperator genotypes are resistant to being cheated on by most defector genotypes from other patches. Such indirect barriers to cheating might be reinforced by unique local patterns of cooperator-defector coevolution (12, 23, 28, 100–102). Avenues for future research include i) investigating the geographic and genetic scale of cooperator-cheater compatibility patches in natural populations and ii) incorporating the potential for cheater-blind barriers to cheating into models of spatially structured social evolution (103), in particular when considering relative contributions of migration vs other forces in shaping equilibrium levels and biogeography of cooperation and cheating.

We propose that limitation of cheating range due to allopatric divergence may be common across diverse social systems. However, allopatrically-evolved forms of kin discrimination (29, 30) are likely to have system-specific rates of evolution and system-specific mechanisms. They may be influenced by the complexity of the cooperative trait being cheated on and the degree of physical proximity between interactants during the social behavior. *M. xanthus* cells produce diverse extracellular compounds during both growth and development (31); there are many possible mechanistic routes by which compatibility of a cooperator and a defector may be reduced, which are unrelated to the mechanism of social defection. Simpler cooperative behaviors, such as production of siderophores that can diffuse to conspecifics at a distance (22), may be less likely to be protected from cheating by nonspecific evolutionary divergence among strains. However, it may still be possible for various siderophores and their receptors (23), and other signal-receptor systems like those for quorum sensing, to diverge in allopatry and thereby generate a biogeographic patchwork of cheating ranges.

## Supporting information

Peer-reviews

## Acknowledgements

We thank Sébastien Wielgoss for assistance with the natural isolate sequence data, Gilles Pütz, and Zachary Blount for productive discussions, and two anonymous reviewers for helpful comments. This study was supported in part by Swiss National Science Foundation (SNSF) grants 31003A/B_16005 to GJV and an ETH Fellowship 16-2 FEL-59 to MV.

## Author contributions

KAS and GJV designed the experiments; KAS, YTNY, and GJV designed the mutants; KAS carried out the experiments; KAS and MV analyzed the data; KAS, YTNY, MV, and GJV wrote the manuscript.

