## Supplemental information for "Allopatric divergence of cooperators confers cheating resistance and limits the effects of a defector mutation"

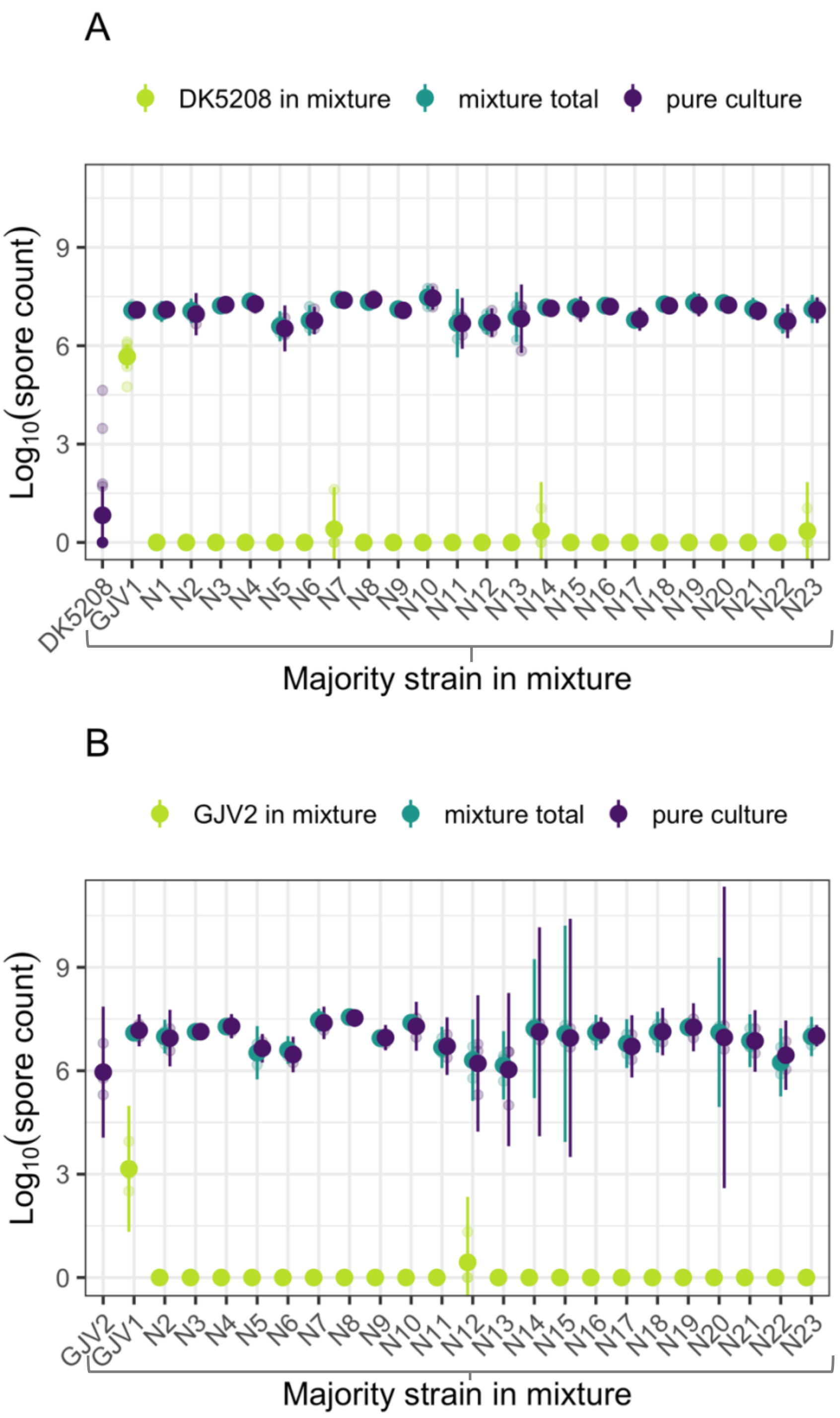


**Fig. S1. Absolute sporulation levels of DK5208, GJV1 and 23 natural isolates in pure culture, DK5208 in mixture with all other strains and total spore production of all mixes.** (A) DK5208 in mixture with GJV1 and 23 natural isolates. (B) GJV2 in mixture with GJV1 and 22 natural isolates. DK5208 and GJV2 started as 1% in all mixed groups. Small circles represent individual replicate estimates, large circles show cross-replicate averages, and error bars show 95% confidence intervals; 3-4 biological replicates for natural isolate mixes, 8 for GJV1.

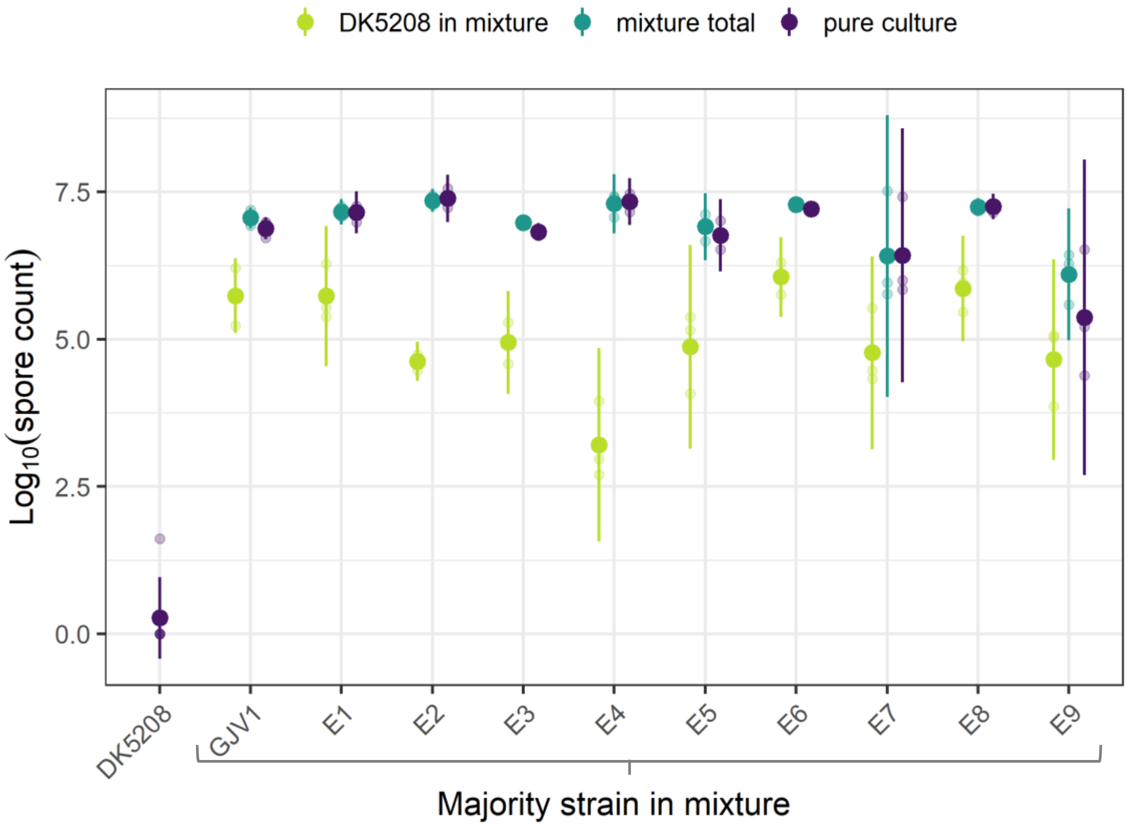


**Fig. S2. Absolute sporulation levels of DK5208, GJV1, and all MyxoEE-3 lab-evolved clones in pure culture, DK5208 in mixture with all other strains, and total spore production of all mixes.** DK5208 started as 1% in all mixed groups. Small circles represent individual replicate estimates, large circles show cross-replicate averages, and error bars show 95% confidence intervals; 3-4 biological replicates.

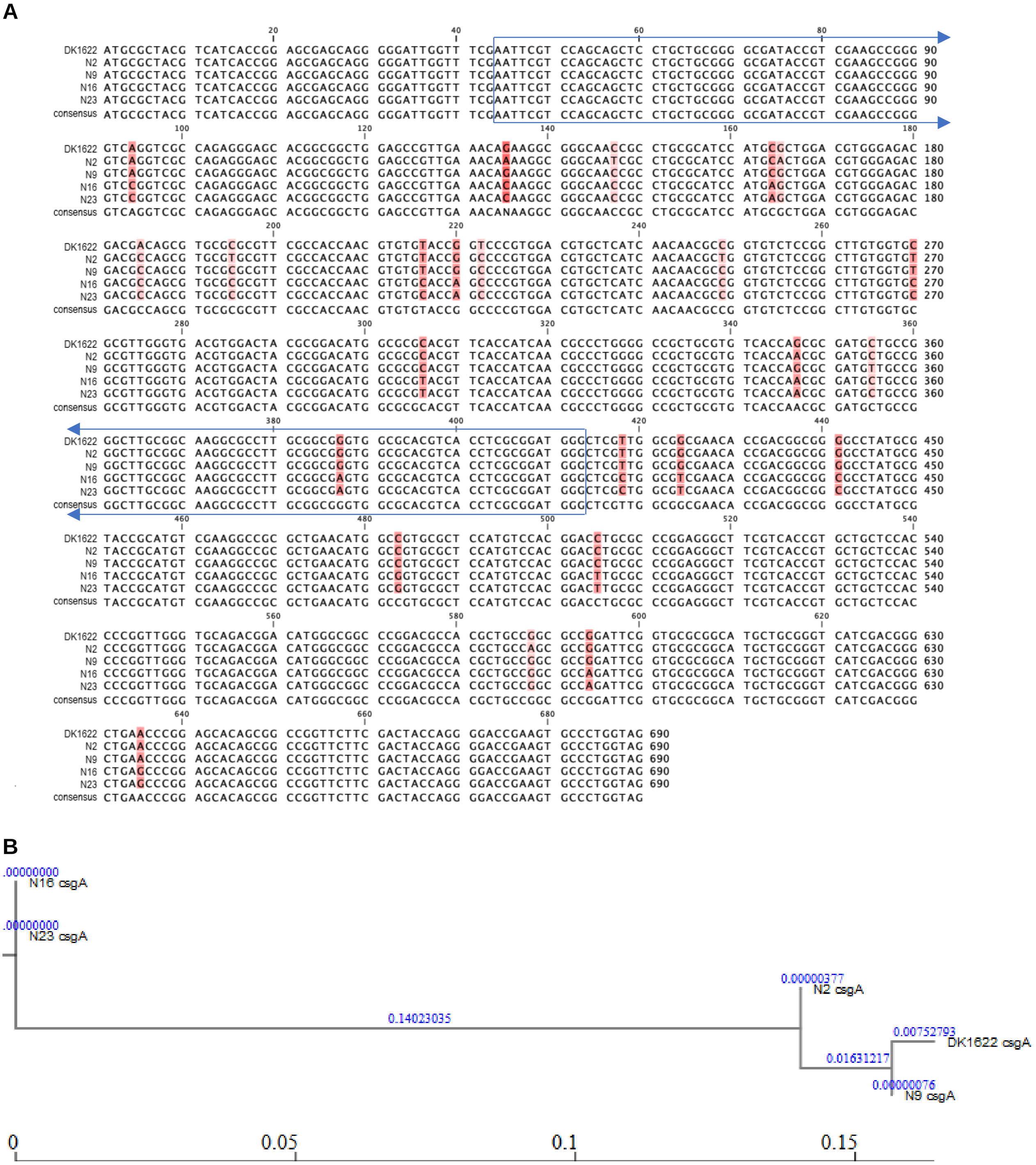


**Fig. S3. *csgA* sequences of *M. xanthus* strains DK1622/GJV1 and four natural isolates disrupted at *csgA* in this study.** (A) Sequence alignment, where red shading indicates polymorphic sites, with darkness reflecting the degree of polymorphism. Boxed regions indicate the beginning and end of the region used as inserts in plasmid construction. (B) Maximum likelihood tree of *csgA* sequences, where blue numbers indicate branch lengths in substitutions per site. The scale bar is in substitutions per site.

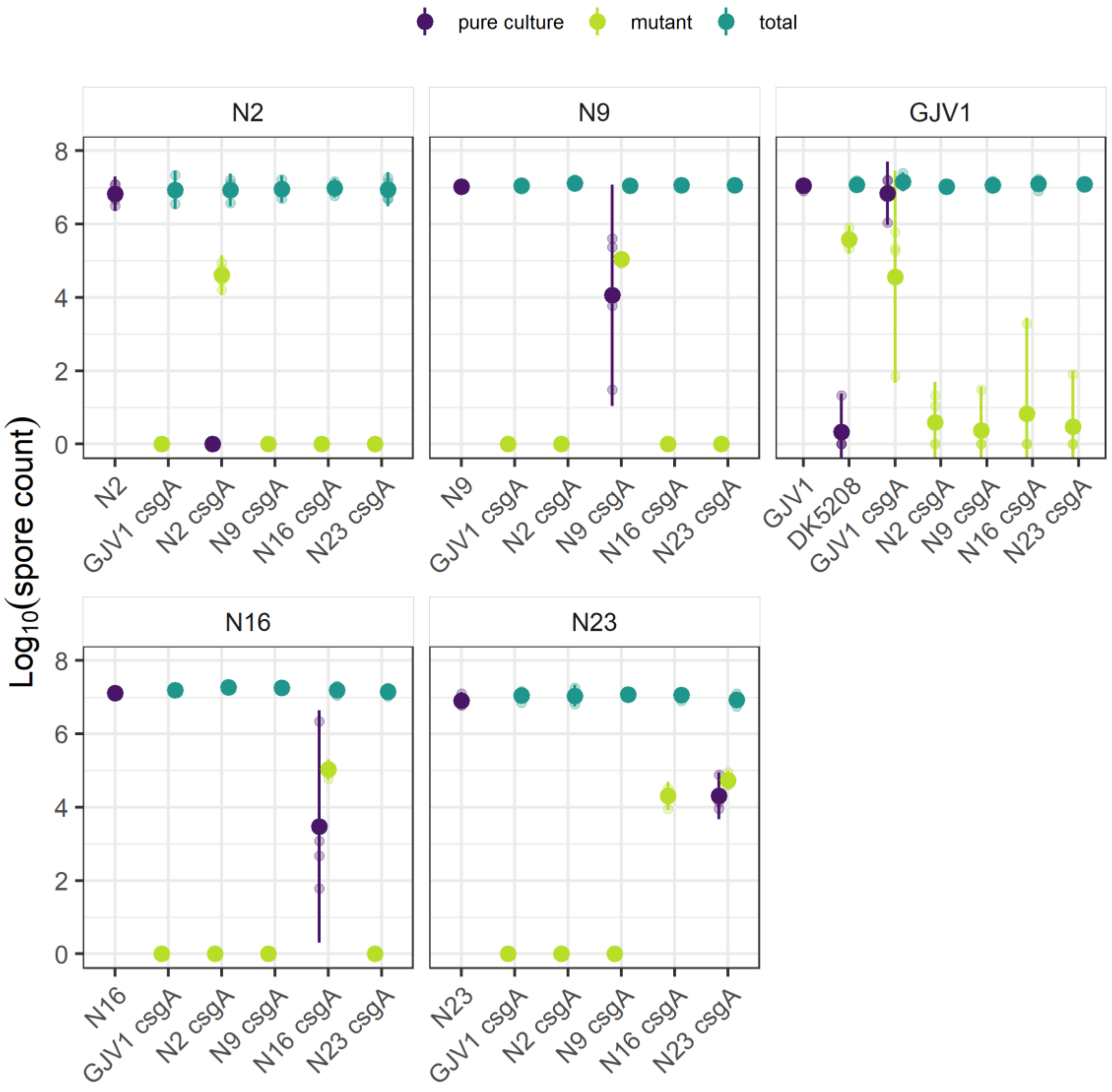


**Fig. S4. Absolute sporulation levels of i) DK5208, GJV1, and four natural isolates in pure culture, ii) each of five new c*sgA* mutants in 1:99 mixture with all five parental strains, and 3) total spore production of all mixes.** Small circles represent individual replicate estimates, large circles show cross-replicate averages, and error bars show 95% confidence intervals; 4 biological replicates.

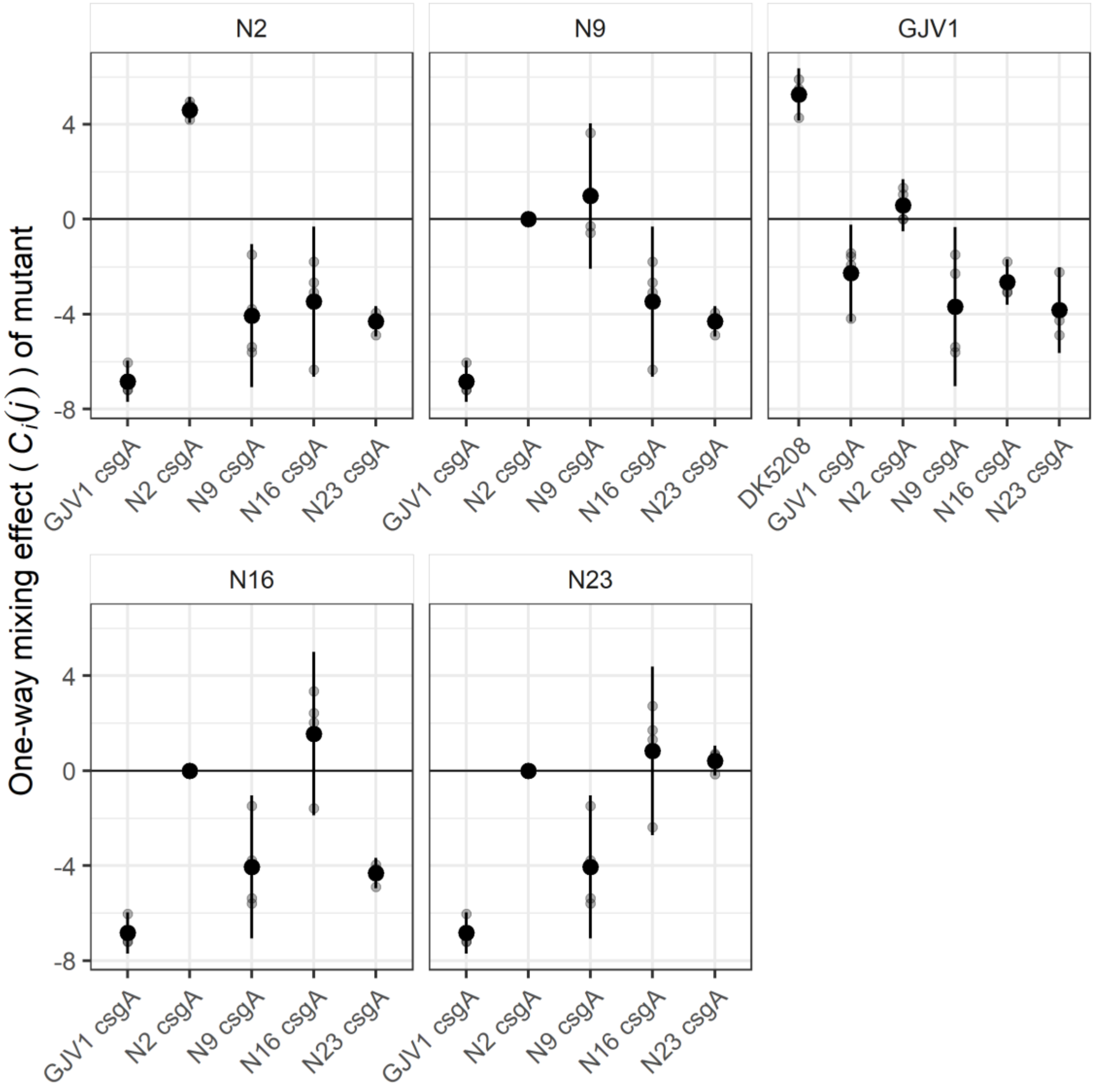


**Fig. S5. One-way mixing effects on *csgA* mutants.** We calculated the degree to which each *csgA* mutant sporulated better or worse in mixture with each proficient strain than it did in pure culture with the parameter $C_i(j)$ (see Methods). Small grey circles represent individual replicate estimates, large black circles show cross-replicate averages, and error bars show 95% confidence intervals; 4 biological replicates.

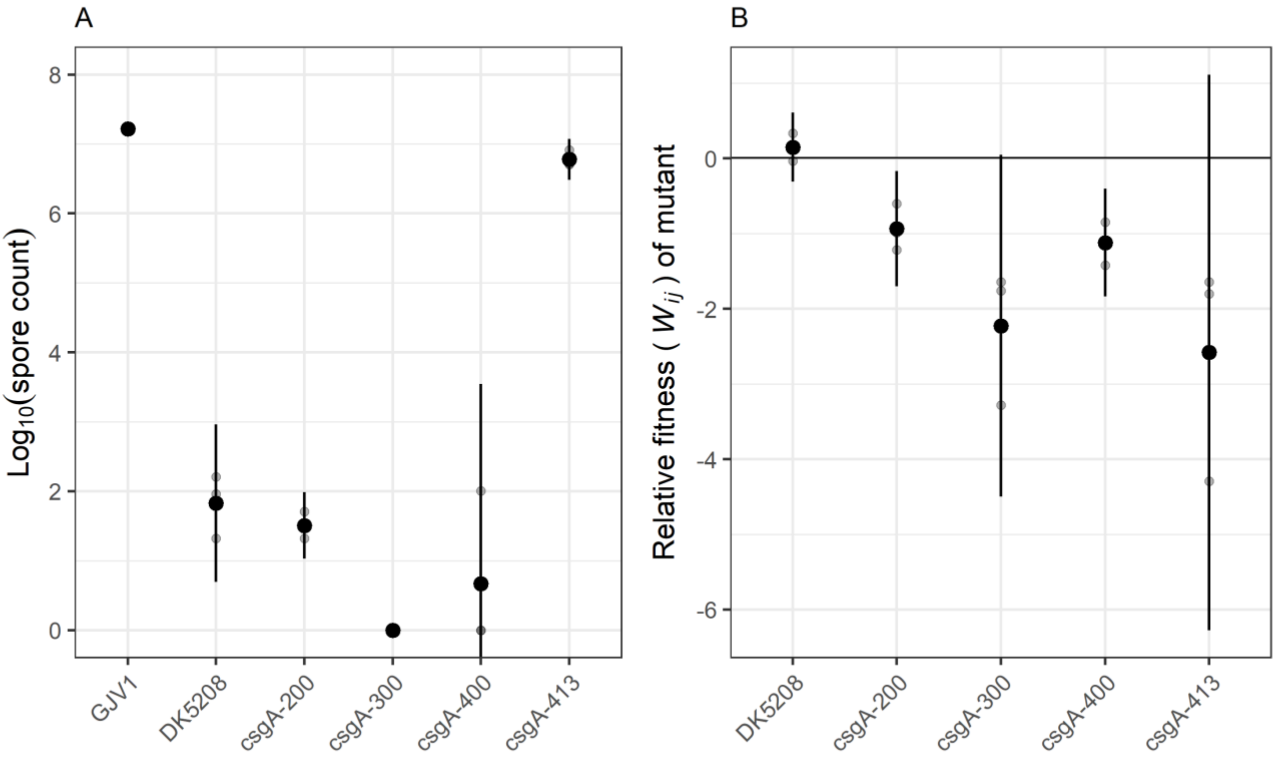


**Fig. S6. Social defects, but no cheating, of different *csgA* mutants in the GJV1 background.** We constructed three additional *csgA* mutants in a similar way to that reported in the Materials and Methods for construction of the csgA-413 mutants. We amplified a 200-bp, 300-bp, and 400-bp fragment of GJV1's native *csgA* allele using one forward primer (5'-TCATCACCGGAGCGAGCAGG-3'; genome positions 1520209-1520228) and three reverse primers, one each for the 200-bp fragment (5'-CACACGTTGGTGGCGAACGC-3'; genome positions 1520394-1520413), the 300-bp fragment (5'-GCCATGTCCGCGTAGTCCAC-3'; genome positions 1520481-1520500), and the 400-bp fragment (5'-CCATCCGCGAGGTGACGTGCGCCA-3'; genome positions 1520587-1520610). We continued with plasmid and mutant construction as reported in the Materials and Methods. We chose these three fragment lengths because we observed that our previous construct, GJV1-csgA (here referred to as csgA-413) had a different pure-culture sporulation phenotype from the existing defector strain DK5208, and we wanted to examine whether this was due to the location of the plasmid insertion in our mutant (413) compared to the transposon insertion in DK5208 (217). We chose fragment lengths that would interrupt *csgA* at positions covering the distance between the two existing mutants. (A) All of the mutants except for csgA-413 are developmentally defective compared to their parent strain GJV1 in mixture ($p$-values < 0.001, csgA-413 $p = 0.9$, post-hoc Tukey HSD test). Strain csgA-300 is more defective than any other mutant except csgA-400 ($p$-values < 0.035, csgA-400 $p = 0.62$, same post-hoc Tukey HSD test). Strain csgA-413 is the least defective of all the mutants ($p$-values < 0.001, same post-hoc Tukey HSD test). (B) All of the mutants constructed in this work show low relative fitness in mixture with GJV1. DK5208 is not statistically shown to cheat in this assay ($p = 0.15$, one-sided t-test for $W_{ij} > 0$). There is slight statistical evidence that the other mutants have reduced fitness relative to GJV1 during this interaction (mean $W_{ij}$ values < 0, 0.05 < $p$-values < 0.1, 4 two-sided t-tests against 0 with Bonferroni-Holm correction). Mutants csgA-300 and csgA-413 have lower relative fitness than does DK5208 ($p$-values < 0.031, post-hoc Tukey HSD test). Given that they are the lowest- and highest-sporulating mutants, respectively, this suggests that the relationship among length of intact csgA gene, social defection phenotype (i.e. amount of spores produced in pure culture), and fitness when mixed with parent (potential for cheating) is complex and nonlinear. Error bars are 95% confidence intervals. Black dots are averages of three biological replicates (grey dots).

**Table S1.** ***M. xanthus*** **strains.** Here we list the original published or new nomenclature of each strain and the simplified nomenclature used in this paper.

| Original or formal name | Ref. | Referred to here as | Genetic manipulation | Resistance | Geographic origin |
|---|---|---|---|---|---|
| DK5208/LS523 | [1, 2] | DK5208 | *csgA*::Tn5-132 | oxytetracycline | [a] |
| GJV1 | [3, 4] | GJV1 | - | - | [a] |
| GJV2 | [3, 5, 6] | GJV2 | spontaneous $rif^R$ mutation | rifampicin | - |
| MyxoEE-3 P02 cycle 40 clone 1 [b] | [7–9] | E1 | - | rifampicin | - |
| MyxoEE-3 P03 cycle 40 clone 1 | [7–9] | E2 | - | - | - |
| MyxoEE-3 P04 cycle 40 clone 1 | [7–9] | E3 | - | rifampicin | - |
| MyxoEE-3 P10 cycle 40 clone 1 | [7–9] | E4 | - | rifampicin | - |
| MyxoEE-3 P12 cycle 40 clone 1 | [7–9] | E5 | - | rifampicin | - |
| MyxoEE-3 P32 cycle 40 clone 1 | [7–9] | E6 | - | rifampicin | - |
| MyxoEE-3 P36 cycle 40 clone 1 | [7–9] | E7 | - | rifampicin | - |
| MyxoEE-3 P38 cycle 40 clone 2 | [7–9] | E8 | - | rifampicin | - |
| MyxoEE-3 P40 cycle 40 clone 1 | [7–9] | E9 | - | rifampicin | - |
| Chihaya 01 | [10] | N1 | - | - | Japan |
| Chihaya 20 | [10] | N2 | - | - | Japan |
| Colombia 01 | [10] | N3 | - | - | Colombia |
| Colombia 03 | [10] | N4 | - | - | Colombia |
| Nei 05 | [10, 11] | N5 | - | - | Mongolia |
| Nei 10 | [10, 11] | N6 | - | - | Mongolia |
| New Jersey 06 | [10, 11] | N7 | - | - | New Jersey, USA |
| New Jersey 10 | [10, 11] | N8 | - | - | New Jersey, USA |
| Serengeti 01 | [10] | N9 | - | - | Tanzania |
| Serengeti 21 | [10] | N10 | - | - | Tanzania |
| Sulawesi 08 | [10, 11] | N11 | - | - | Indonesia |
| Tubingen C22 | [10, 11] | N12 | - | - | Germany |
| Tubingen C42 | [10, 11] | N13 | - | - | Germany |
| GH2.1.4c40 | [12] | N14 | - | - | Indiana, USA (“Greg’s House”) |
| GH3.2.7C | [12] | N15 | - | - | Indiana, USA (“Greg’s House”) |
| GH3.5.6c2 | [12] | N16 | - | - | Indiana, USA (“Greg’s House”) |

| | | | | | |
|---|---|---|---|---|---|
| GH5.1.9c20 | [12] | N17 | - | - | Indiana, USA (“Greg’s House”) |
| KF2.1.1B | [12] | N18 | - | - | Indiana, USA (“Kent Farms”) |
| KF3.2.8c11 | [12] | N19 | - | - | Indiana, USA (“Kent Farms”) |
| KF5.4.6c29 | [12] | N20 | - | - | Indiana, USA (“Kent Farms”) |
| MC3.1.9c3 | [12] | N21 | - | - | Indiana, USA (“Moore’s Creek”) |
| MC3.2.6B | [12] | N22 | - | - | Indiana, USA (“Moore’s Creek”) |
| MC3.5.9c15 | [12] | N23 | - | - | Indiana, USA (“Moore’s Creek”) |
| GJV1_*csgA483* | this work | - | *csgA*::pCR-*csgA483* | kanamycin | - |
| Chihaya 20_*csgA483* | this work | N2 *csgA* | *csgA*::pCR-*csgA483* | kanamycin | - |
| Serengeti 01_*csgA483* | this work | N9 *csgA* | *csgA*::pCR-*csgA483* | kanamycin | - |
| GH3.5.6c2_*csgA483* | this work | N16 *csgA* | *csgA*::pCR-*csgA483* | kanamycin | - |
| MC3.5.9c15_*csgA483* | this work | N23 *csgA* | *csgA*::pCR-*csgA483* | kanamycin | - |

[a]Strain DK1622 – the lab progenitor of both DK5208 and GJV1 – is reported to be derived from a strain isolated in Ames, Iowa, USA [13].
[b]Multiple clones were selected and stored frozen after MyxoEE-3 cycle 40.

**Table S2. History and mutations of MyxoEE-3 lab-evolved strains.** The strains referred to here as E1-E9 are clones from populations of *M. xanthus* that evolved as vegetatively growing colonies swarming across hard (1.5%) or soft (0.5%) agar [7, 8] over 40 two-week growth cycles. Here we provide information about the evolution conditions and accumulated mutations of each strain (from Table S4 in [7]).

| Strain | Original name | Agar concentration | Mutations | Genic | Coding | Synonymous | Intergenic |
|---|---|---|---|---|---|---|---|
| E1 | P02 clone 1 | 1.5% | 13 | 10 | 8 | 2 | 3 |
| E2 | P03 clone 1 | 1.5% | 10 | 10 | 10 | 0 | 0 |
| E3 | P04 clone 1 | 1.5% | 10 | 9 | 9 | 0 | 1 |
| E4 | P10 clone 1 | 1.5% | 10 | 10 | 7 | 3 | 0 |
| E5 | P12 clone 1 | 1.5% | 19 | 15 | 14 | 1 | 4 |
| E6 | P32 clone 1 | 0.5% | 12 | 12 | 11 | 1 | 0 |
| E7 | P36 clone 1 | 0.5% | 14 | 14 | 13 | 1 | 0 |
| E8 | P38 clone 2 | 0.5% | 10 | 10 | 9 | 1 | 0 |
| E9 | P40 clone 1 | 0.5% | 13 | 12 | 11 | 1 | 1 |

**Table S3. Plasmids constructed in this work.**

| Plasmid name | *csgA* allele |
|---|---|
| pGJV1*csgA483* | GJV1 |
| pChihaya20*csgA483* | Chihaya 20 |
| pSerengeti01*csgA483* | Serengeti 01 |
| pGH356c2*csgA483* | GH3.5.6c2 |
| pMC359c15*csgA483* | MC3.5.9c15 |
