## Supplementary material for "Allopatric divergence of cooperators confers cheating resistance and limits the effects of a defector mutation": Peer-reviews

Reviews of manuscript version 1

Editorial Board Member's Remarks to Author:

This is an interesting paper of potential interest to readers. Yet the manuscript could be improved as indicated by the referees. Besides these specific comments, the likely generality or lack thereof of the result should be made more clear.

Reviewer Comments:

Reviewer #1:

Suitable Quality?: Yes
Sufficient General Interest?: Yes
Conclusions Justified?: Yes
Clearly Written?: Yes
Procedures Described?: Yes
Supplemental Material Warranted?: Yes

Comments:
Schaal et al. test the extent to which the fitness effects of a mutation are influenced by social context and by genomic background in *Myxococcus xanthus*. Specifically, they look at a mutation in a gene that produces signals for fruiting body formation, *csgA*. Disruption of this gene allows a lab strain to cheat on a close relative by decreasing the cheat's investment in the fruiting body and increasing its representation in the spore forming cells. First, the authors demonstrate that this mutant cannot cheat on wild isolates. They then show that experimental evolution of the lab cooperator can decrease the ability of the mutant to cheat. Finally, the authors look at the impact of moving the mutation that causes cheating in the lab strain into wild isolates. They find that the mutation has different impacts on spore formation in different backgrounds. In the cases that the mutation disrupts spore formation, the mutant can be rescued by its parent strain but rarely by other wild isolates. The authors marshal this data to suggest that cooperation can be maintained by genetic diversification, and that the genetic basis of a conserved developmental phenotype can diverge across lineages.

In the experiments testing the lab cheater against other genotypes, it is unclear to me whether results are specific to scenarios of cooperation. In some sense if you have any genotype A that is more fit than a genotype B, is it necessarily surprising that A might be less fit than genotypes C to F? Similarly, if you allow genotype B to evolve wouldn't you expect that it will often become more fit than A? It seems like it would help to disentangle social interactions versus other aspects of the genotypic differences by knowing the fitness of GJV1 (the lab cooperator) against the wild isolates and the evolved isolates.

It would also be helpful to see more discussion of the generality of these findings. It seems likely that the extent to which divergence protects against cheaters will be influenced by the nature of cooperation. For example, secretion of public goods may be less protected by divergence than developmental phenotypes. It would also be good to see some more discussion of how the current results relate to previous work on the rapid evolution of cooperative signals (e.g. PMID 16572169). Some work on evolution of signals is cited but there is little discussion of its relation to the current work.

This is a well-written paper and social evolution and evo-devo are certainly of broad interest. It is perhaps somewhat incremental building on some of the senior authors' previous work showing that kin discrimination can rapidly evolve in this system (PMID 26150498).

Minor comment:
Line 176 - The incorrect figure is cited.

Line 178 - It would be interesting to know how the difference in outcome compares to the genetic similarity. Sequences are provided in Figure S3, but it would be helpful to see this data converted to a tree. Similarly, I would be interested to know the similarity of the broader genetic background. Is this information known?

Reviewer #2:

Suitable Quality?: Yes
Sufficient General Interest?: Yes
Conclusions Justified?: Yes
Clearly Written?: Yes
Procedures Described?: Yes
Supplemental Material Warranted?: Yes

Comments:
In this manuscript by Schaal et al., the authors present data that show cheating effects in *Myxococcus xanthus* are limited to closely related cooperator-cheater pairs and a relatively small genetic divergence can alter the outcome of cooperator-cheater interactions. The authors also find that a cheating mutation in the *csgA* gene confers a cheating interaction in one genetic background but not in others. The authors argue that this inability of this *csgA* mutation to recapitulate cheating in other strain backgrounds is an example of Developmental System Drift in a microbial system, as *csgA* is conserved between strains but seems to have different functions in spore production. In general, I thought the manuscript was well written and clear, although there was some jargon that could be better defined. My only concerns are to what extent other interactions such as competitive antagonisms might be masking cheating interactions, which I've outlined below.

1) The authors state that antagonisms between *M. xanthus* strains are commonly found, and my concern is to what extent is the observed pattern in Figure 1 inhibition by the natural strains vs. the inability of DK5208 to be complemented by the natural strains? I am not familiar when competitive inhibition between strains occurs in *M. xanthus* so this may not be an issue during development. However, if there is some form of inhibition that should be visible when competing the ancestral, cooperative DK1622 strain against the natural strains. Is this strain still able to form spores? However, with that said, I found Figure 2 much more convincing, as it is unlikely that any inhibition between recently diverged stains would have evolved.

Similarly, in Figure 4, to what extent is it the *csgA* mutation is determining the fitness outcomes observed? For example, in panel one when GJV1 without the mutation (or N9, N16, N23) is competed against N2 is it able to produce spores?

This seems to be important as there is presumably a difference between the inability of a cheating strain to be complemented or being actively inhibited via some sort of toxic intermediary.

2) On line 40 in the abstract, it states that genomic divergence determined whether and how much the same mutation reduces a cooperative phenotype. Based on Figure 3, I'm confused by this statement. To what extent is genomic divergence predictive of the effects of the *csgA* mutation?

3) The authors used the term 'cheater-blind' throughout the manuscript without clearly explaining what they mean by this. It would have been helpful if this was defined in the abstract or at the very least in the introduction.

4) On line 200, the manuscript states that the mutants socially exploited their parents. Is it exploitation if there is no fitness consequence on the parental strain?

5) I found the title unclear and confusing. Neither the latency of cheater evolution, nor the genetic requirements were explicitly measured in these experiments. It may very well be that because high cheater loads in wild populations of *M. xanthus* there are strong pre-adaptations or genetic architecture that predisposes diversification of cheatable developmental pathways. This isn't discussed, so again to what extent is this evolution really latent? Also, what are the genetic requirements for a cooperative trait in this system? This isn't explicitly addressed apart from suggesting there are genetic background effects, but that's not necessarily informative about what the genetic requirements are.

Response letter January 14, 2021

We thank the reviewers for their positive, thoughtful comments and for bringing these points to our attention. We have addressed them as follows.

Reviewer Comments:

Reviewer #1
Comments:

*In the experiments testing the lab cheater against other genotypes, it is unclear to me whether results are specific to scenarios of cooperation. In some sense if you have any genotype A that is more fit than a genotype B, is it necessarily surprising that A might be less fit than genotypes C to F? Similarly, if you allow genotype B to evolve wouldn't you expect that it will often become more fit than A? It seems like it would help to disentangle social interactions versus other aspects of the genotypic differences by knowing the fitness of GJV1 (the lab cooperator) against the wild isolates and the evolved isolates.*
We have added a figure (new Fig. S1B) showing that a developmentally proficient marked variant of the GJV1 wild-type (rifampicin-resistant GJV2) fails to sporulate when mixed as the minority strain with natural isolates.

We have no reason to think that the genomic differences causing the fitness of the cheater DK5208 to be lower in mixture with natural isolates or lab-evolved strains than in mixture with GJV1 are specific to C-signalling or any other development-specific genes (assuming that is what the reviewer means by "specific to scenarios of cooperation"). For the natural isolate mixes, we expect that toxins produced by the natural isolates are responsible for their nearly complete inhibition of sporulation by the cheater DK5208 – toxins that are also generally expressed during vegetative growth (Rendueles et al. 2015, *Current Biology*). Consistent with this, like the cheater DK5208, the marked variant of GJV1 (GJV2) fails almost entirely to make any spores in mixture with any of the natural isolates examined here. Analogous experiments mixing GJV2 with the lab-evolved lines were not performed because all but one of those lines are lab descendants of GJV2 and thus also resistant to rifampicin. However, as we note in the manuscript, because we know in detail the evolutionary history of the lab-evolved lines since they diverged from GJV1, we can unequivocally state that the divergence between these lines and the cheater is not due to any selection on fitness during development, much less on developmental interactions involving C-signal specifically. Also, we now refer to genome sequences of several of the lab-evolved lines, including the two that most strongly prevent cheating by DK5208 (E2 and E4), in which there were no mutations in *csgA.*

*It would also be helpful to see more discussion of the generality of these findings. It seems likely that the extent to which divergence protects against cheaters will be influenced by the nature of cooperation. For example, secretion of public goods may be less protected by divergence than developmental phenotypes. It would also be good to see some more discussion of how the current results relate to previous work on the rapid evolution of cooperative signals (e.g. PMID 16572169). Some work on evolution of signals is cited but there is little discussion of its relation to the current work.*
We expand on the discussion of how these findings may relate to other cooperative systems in order to make the level of generality clearer, specifically as regards different types of cooperative behaviors and their relative complexities. We also emphasize more the theme of signal evolution but opt to not venture into discussing potential scenarios involving greenbeards specifically.

*Minor comment:*
*Line 176 - The incorrect figure is cited.*
We corrected this.

*Line 178 - It would be interesting to know how the difference in outcome compares to the genetic similarity. Sequences are provided in Figure S3, but it would be helpful to see this data converted to a tree.*
We provide such a tree as a new panel in Figure S3. We find that there is no correlation across strains between the difference in the magnitude of *csgA*-disruption effects shown in Fig 3 and degree of divergence at *csgA*. For example, the *csgA* sequences of GJV1 and N2 are relatively similar but these two strains show the greatest difference in the magnitude of the *csgA*-disruption effect. In contrast, N9 and N23 have very similar average *csgA*-disruption effects (albeit with much variation about the N9 mean) while having highly divergent *csgA* sequences.

*Similarly, I would be interested to know the similarity of the broader genetic background. Is this information known?*
Divergence among four of the five parents of the mutants featured in Fig. 3 based on whole-genome sequences is shown in Fig. 1 of Rajagopalan *et al.* 2015. (The only parent strain not included in Rajagopalan et al. is GH3.5.6c2. Divergence between this strain, MC3.5.9c15 and DK1622 based on a concatemer of conserved genes can be seen in Fig. S2 of Kraemer and Velicer 2011.) As is the case for the *csgA* sequences noted above, there is clearly no correlation across strains between the difference in the magnitude of *csgA*-disruption effects shown in Fig 3 and the degree of overall genomic divergence.

Reviewer #2
Comments:

*1) The authors state that antagonisms between M. xanthus strains are commonly found, and my concern is to what extent is the observed pattern in Figure 1 inhibition by the natural strains vs. the inability of DK5208 to be complemented by the natural strains? I am not familiar when competitive inhibition between strains occurs in M. xanthus so this may not be an issue during development. However, if there is some form of inhibition that should be visible when competing the ancestral, cooperative DK1622 strain against the natural strains. Is this strain still able to form spores? However, with that said, I found Figure 2 much more convincing, as it is unlikely that any inhibition between recently diverged stains would have evolved.*
*Similarly, in Figure 4, to what extent is it the csgA mutation is determining the fitness outcomes observed? For example, in panel one when GJV1 without the mutation (or N9, N16, N23) is competed against N2 is it able to produce spores? This seems to be important as there is presumably a difference between the inability of a cheating strain to be complemented or being actively inhibited via some sort of toxic intermediary.*

All evidence suggests that the non-competitiveness of the cheater vs. natural isolates is due to generic inhibition by the natural strains rather than any mechanism specific to interacting with a cheater. We addressed these questions collectively in lines 213-220 of our first submission while discussing the results shown in Fig. S4:
"Like DK5208 and the natural isolate *csgA* mutants, GJV1 *csgA* failed to sporulate effectively in mixtures with natural isolates (Fig. S4). Because GJV1 *csgA* exhibits no pure-culture sporulation

defect, its total suppression by four natural isolates indicates that these strains generically antagonize GJV1/DK1622-background strains. Thus, **the inability of DK5208 to sporulate in the presence of natural isolates (Fig. 1) appears to be due to generic antagonisms to which all GJV1/DK1622-background strains are susceptible** rather than to either a mechanistic inability of the DK5208 *csgA* defect to be complemented by the CsgA proteins of natural isolates or antagonistic behavior directed specifically at cheater genotypes."

In response to this comment and a comment by Reviewer 1 on the same theme, we now emphasize this point by adding a new supplementary figure showing that a developmentally proficient, marked variant of GJV1 with no mutation in *csgA* (GJV2) is also strongly inhibited by the natural isolates. We now also mention that antagonisms between natural isolates, which are the norm in pairwise competition experiments (see Rendueles et al. 2015, *Current Biology*), appear to generally be manifested during both vegetative growth and development, further showing their non-specificity toward cheaters.

*2) On line 40 in the abstract, it states that genomic divergence determined whether and how much the same mutation reduces a cooperative phenotype. Based on Figure 3, I'm confused by this statement. To what extent is genomic divergence predictive of the effects of the csgA mutation?*
We did not intend to connote that the degree to which genomes have diverged correlates with the degree to which the focal mutation reduces the phenotype. We rather meant simply that particularities of the *overall character* of divergence – whether degree of divergence, type of divergence at particular loci, or both – determine this. We rephrased the line in question so as not to indicate a correlative relationship.

*3) The authors used the term 'cheater-blind' throughout the manuscript without clearly explaining what they mean by this. It would have been helpful if this was defined in the abstract or at the very least in the introduction.*
We added a clarifying phrase to the abstract and a definition of the term in the Introduction.

*4) On line 200, the manuscript states that the mutants socially exploited their parents. Is it exploitation if there is no fitness consequence on the parental strain?*
Yes, we follow precedent of 'exploitation' being defined simply as using an available resource, opportunity or interaction to one's benefit – in our specific case as one strain gaining in absolute fitness from interacting with another, irrespective of the effect of the interaction on the latter (e.g. Kraemer & Velicer 2014, Proc B). Here the term is defined solely in terms of the positive effect of the interaction on the recipient and thus has no inherent negative connotation regarding the other interactant, even though such negative effects do often occur. We added a statement to the Semantics section of the Methods to clarify this.

*5) I found the title unclear and confusing. Neither the latency of cheater evolution, nor the genetic requirements were explicitly measured in these experiments. It may very well be that because high cheater loads in wild populations of M. xanthus there are strong pre-adaptations or genetic architecture that predisposes diversification of cheatable developmental pathways. This isn't discussed, so again to what extent is this evolution really latent?*
We altered the title for greater clarity. Throughout the first submission we used 'latent' (and variants thereof) to refer to evolutionary processes in which the genetic basis of a phenotype - e.g. resistance to being cheated on - evolves in a hidden manner by any evolutionary mechanism other than direct selection for the phenotype, which is only manifested later in a

different context (e.g. see line 161 of our first submission). In our study, for example, we know that the resistance of lab-evolved clones E2, E4 and others to being cheated on by DK5208 evolved latently, i.e. during an evolution experiment in which lineages leading to those clones not only never encountered DK5208, but never even underwent development at all. We opt to continue using this term throughout the manuscript in this sense, but agree to remove it from the title to avoid potential confusion before readers can see the sense in which we use the term. The title now refers to allopatric divergence, which implies latent evolution of interaction phenotypes between lineages that had never previously met. We qualify use of 'allopatric divergence' with respect to the natural isolates in the semantics section.

*Also, what are the genetic requirements for a cooperative trait in this system? This isn't explicitly addressed apart from suggesting there are genetic background effects, but that's not necessarily informative about what the genetic requirements are.*

By 'genetic requirements' we were referring to the length of uninterrupted *csgA* required to generate the cooperation-based trait of a high level of spore production. See lines 178-181 of our first submission. In our revision we have added a new sentence to the end of the same paragraph to emphasize this point even more using the same terminology employed in the title.
